# Incoherent feedforward regulation via Sox9 and Erk underpins mouse tracheal cartilage development

**DOI:** 10.1101/2020.07.22.216747

**Authors:** Takuya Yoshida, Michiyuki Matsuda, Tsuyoshi Hirashima

## Abstract

Tracheal cartilage provides architectural integrity to the respiratory airway, and defects in this structure during embryonic development cause severe congenital anomalies. Previous genetic studies have revealed genes that are critical for the development of tracheal cartilage. However, it is still unclear how crosstalk between these proteins regulates tracheal cartilage formation. Here we show a core regulatory network underlying murine tracheal chondrogenesis from embryonic day (E) 12.5 to E15.5, by combining volumetric imaging of fluorescence reporters, inhibitor assays, and mathematical modeling. We focused on SRY-box transcription factor 9 (Sox9) and extracellular signal-regulated kinase (Erk) in the tracheal mesenchyme, and observed a synchronous, inverted U-shaped temporal change in both Sox9 expression and Erk activity with a peak at E14.5, whereas the expression level of downstream cartilage matrix genes, such as collagen II alpha 1 (*Col2a1*) and aggrecan (*Agc1*), monotonically increased. Inhibitor assays revealed that the Erk signaling pathway functions as an inhibitory regulator of tracheal cartilage differentiation during this period. These results suggest that expression of the cartilage matrix genes is controlled by an incoherent feedforward loop via Sox9 and Erk, which is supported by a mathematical model. Furthermore, the modeling analysis suggests that a Sox9-Erk incoherent feedforward regulation augment the robustness against the variation of upstream factors. The present study provides a better understanding of the regulatory network underlying the tracheal development and will be helpful for efficient induction of tracheal organoids.

## Introduction

The mammalian trachea is a tubular organ of the respiratory system and is composed of several tissues from different origins, such as endoderm-derived epithelium and mesoderm-derived cartilage (Cardoso and Lü, 2006). The tracheal cartilage, also known as the tracheal ring, exhibits a C-shaped semi-ring architecture that surrounds the epithelial airway on the ventral side and provides structural support. Abnormal formation of the tracheal rings can collapse the airways and obstruct breathing, which leads to congenital defects, including tracheomalacia and tracheal stenosis (Arooj Sher and J Liu, 2016). Thus, a fundamental understanding of the processes underpinning tracheal ring development is essential, yet it is still incomplete.

The development of tracheal rings has been investigated using mouse genetics, which has revealed the importance of multiple transcription factors and signaling pathways. Among these, SRY-box transcription factor 9 (SOX9) is known to be a master regulator that plays a critical role in cartilage differentiation by inducing gene expression of key cartilage matrix molecules, such as collagen II alpha 1 (Col2a1), and aggrecan (Agc1) (Bi et al., 1999; Han and Lefebvre, 2008). It has been demonstrated that SOX9 functions in each successive step of the cartilage differentiation processes, including mesenchymal condensation, commitment to the chondroprogenitor, and maintenance of proliferating chondrocytes (Bi et al., 1999; Akiyama et al., 2002). The importance of Sox9 in tracheal development was revealed by reports which described a complete absence of the tracheal rings in mesenchymal *Sox9* knockout mice (Hines et al., 2013; Turcatel et al., 2013). In addition, the haploinsufficiency of *Sox9* caused hypoplastic cartilage formation, indicating that Sox9 dosage is critical to tracheal ring formation (Bi et al., 2001). In sonic hedgehog (Shh) knockout mice, *Sox9* mRNA expression was lost in the developing tracheae at a later stage of tracheal ring development, leading to the failure of tracheal ring formation, suggesting that Shh plays an important role in controlling Sox9 expression (Park et al., 2010).

Another important signaling pathway is the fibroblast growth factor (Fgf) – extracellular signal-regulated kinase (Erk) signaling axis. A previous study showed that severe malformations of the tracheal rings were observed when *Fgf10* or its main receptor *Fgfr2b* were either ubiquitously knocked out or overexpressed in the tracheal mesenchyme. These results suggest that FGF10 expression levels must be within a specific range to ensure correct formation of the tracheal rings (Tiozzo et al., 2009; Sala et al., 2011). The most downstream kinase of the signaling cascade, ERK, is considered to be essential for tracheal development because the genetic deletion of both *Mek1* and *Mek2* (upstream kinase of ERK) in the mesenchyme resulted in defective tracheal rings (Boucherat et al., 2015). These studies make a strong case for the necessity of Fgf-Erk signaling in the normal development of the tracheal rings; however, the mechanisms through which ERK activation regulates cartilage differentiation are still unknown. It is noteworthy that there are several studies which have presented conflicting results; on one hand, a positive role for Erk signaling was shown using mouse primary chondrocytes (Murakami et al., 2000), while on the other, a negative role was described in chicken embryonic limb buds (Oh et al., 2000; Bobick and Kulyk, 2004).

In this study, we explore the crosstalk between the Shh-Sox9 and Fgf-Erk signaling pathways and the regulatory network that contributes to tracheal ring development. We first show a synchronized temporal profile of Sox9 expression and Erk activity using volumetric imaging of fluorescence reporters. Combined with inhibitor assays, we then show that cartilage matrix genes are positively regulated by Sox9, and in parallel, negatively regulated by Erk activity, in an incoherent feedforward manner. Finally, a mathematical model demonstrates that an incoherent feedforward loop via Sox9 and Erk can explain the dynamics of cartilage matrix gene expression during tracheal cartilage formation.

## Results

### Mesenchymal condensation begins between E12.5 and E13.5

To morphologically characterize the developing murine trachea, we dissected tracheae from E12.5 to E14.5 mouse embryos and processed the tissues using whole mount immunohistochemistry to examine the staining of the nuclei and plasma membrane of the tracheal epithelium. We focused on the ventral region of the tracheal mesenchyme, where the tracheal cartilage rings are formed, and we show the frontal and sagittal planes of the ventral region (Fig. 1A). We also show the flattening of nuclei, which were used to evaluate how flat the best fitting ellipse should be in comparison to a circle for nuclei evaluations (Fig. 1B).

**Figure 1.**
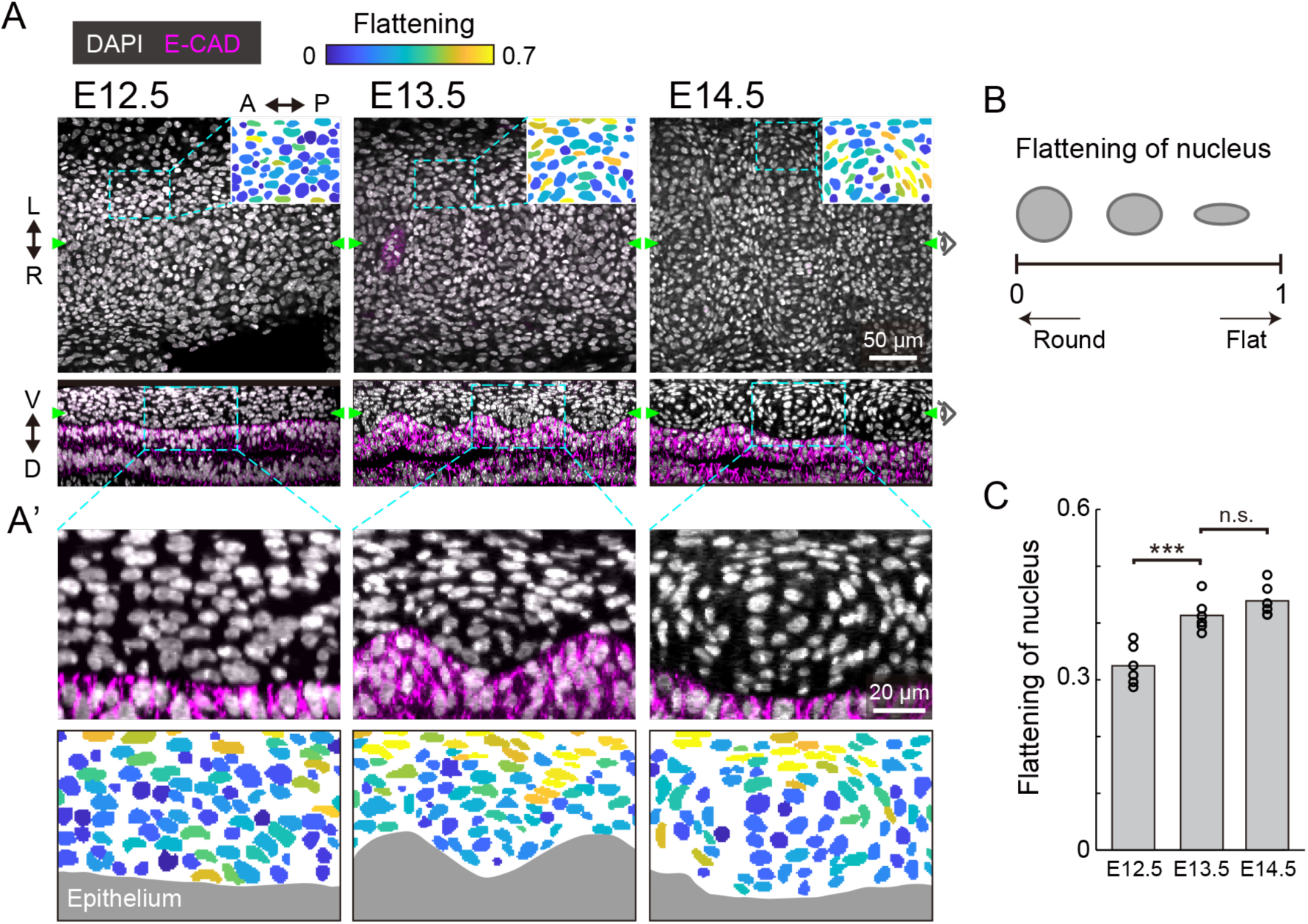
Nuclear distribution in the developing trachea. (A) Immunofluorescence section images of anti-E-cadherin (magenta) with DAPI nuclear counterstaining (white) in developing murine tracheae from E12.5 to E14.5 (upper: frontal plane, lower: sagittal plane). In the pictographic transcriptions in the top right corner, color denotes the flattening of nuclei seen in the dotted windows. Green arrowheads indicate the point visualized in the orthogonal views. Scale bar, 50 µm. (A’) Magnified images corresponding to the dotted windows in the lower row of (A) (upper) and the flattening of nuclei. Gray indicates epithelium. Scale bar, 20 µm. L: left, R: right, A: anterior, P: posterior, V: ventral, D: dorsal. (B) Schematics of the flattened nuclei. (C) Flattening of nucleus from E12.5 to E14.5 in the sagittal plane. Each point represents the mean value within the sample. Welch’s two-sample t-test, E12.5-E13.5: p<0.001. E13.5-E14.5: p=0.152. n=6.

At E12.5, the mesenchymal nuclei were almost round and homogeneously distributed, in both the frontal (Fig. 1A) and the sagittal planes (Fig. 1A’). At E13.5, some mesenchymal nuclei were more elongated and surrounded the cells close to the epithelium, forming a template of the chondrogenic nodule (Fig. 1A and 1C). This process is known as mesenchymal condensation, the initial step of chondrogenesis in general (Goldring et al., 2006). These condensations were observed between the ridges of wavy tracheal epithelium (Fig. 1A’). At E14.5, the peripheral nuclei of the mesenchymal condensations were further elongated and the azimuthal polarization pattern was observed (Fig. 1A-1C). This phenomenon is consistent with the formation of the perichondrium, the next step of mesenchymal condensation during chondrogenesis (Goldring et al., 2006). These observations indicate that mesenchymal cell differentiation into chondrocytes of the tracheal cartilage begins between E12.5 and E13.5.

### Sox9 expression increases up to E14.5, and decreases thereafter

We next examined the expression of Sox9, the early differentiation marker of chondrocytes, in the developing murine tracheae from E12.5 to E15.5. For this purpose, we employed Sox9-EGFP knock-in mice, in which the expression level of EGFP has been shown to correlate with the endogenous expression of Sox9 (Nel-Themaat et al., 2009; Nakamura et al., 2011). EGFP fluorescence in the ventral side of the epithelium was observed using two-photon microscopy.

Observation by 3D imaging showed the appearance of distinct cell clusters, defined by EGFP intensity, from E13.5 onward (Fig. 2A). These high-EGFP clusters are located between epithelial ridges, corresponding to the mesenchymal condensations (Fig. 1A and Fig. S1A). The condensed mesenchymal cells with high EGFP intensity also exhibited a C-shaped semi-ring structure, indicating formation of the chondrogenic nodule (Fig. S1B). We then quantified the EGFP intensity in each mesenchymal cell. Although EGFP expression was uniform along the antero-posterior axis in the median line at E12.5, the periodic pattern of EGFP intensity was confirmed at E13.5 and the amplitude of this periodic intensity became larger at E14.5 (Fig. 2B). The median EGFP intensity in the mesenchymal condensations increased 1.18-fold from E12.5 to E13.5, and 1.65-fold from E13.5 to E14.5 (Fig. 2C), which linearly correlates with Sox9 protein levels (Fig. S1C and S1D). However, the EGFP intensity in the mesenchymal condensations decreased 0.56-fold from E14.5 to E15.5 (Fig. 2A-2C), indicating that Sox9 expression is suppressed at E14.5.

**Figure 2.**
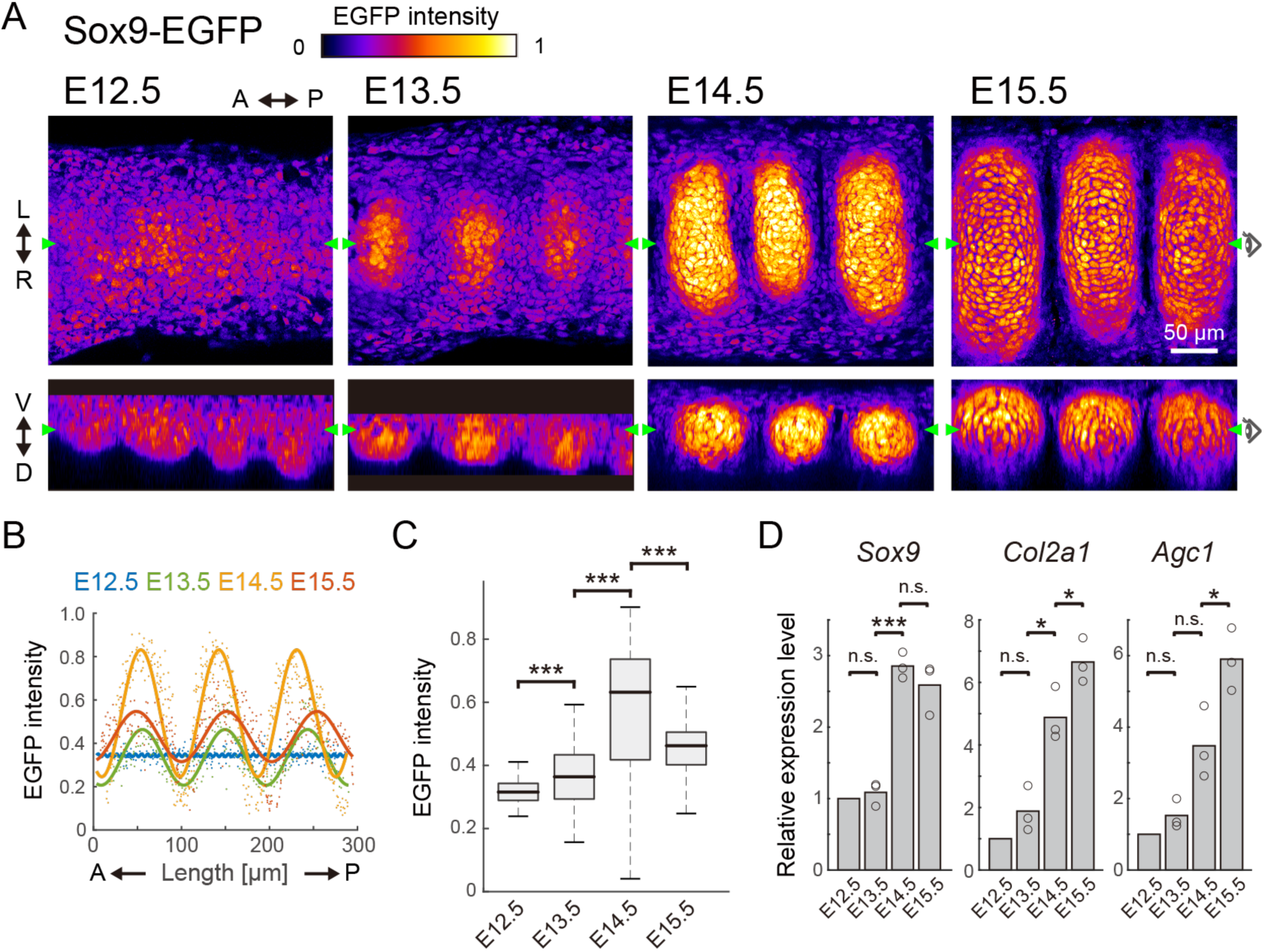
Sox9 expression profile in the developing trachea. (A) Section view of EGFP in the frontal (upper) and sagittal (bottom) planes from E12.5 to E15.5. Color denotes the normalized EGFP intensity. Green triangles in each plane represent the position of orthogonal sections. Scale bar, 50 µm. L: left, R: right, A: anterior, P: posterior, V: ventral, D: dorsal. (B) EGFP intensity in each mesenchymal cell along the antero-posterior axis. Dots represent mean intensity in each cell and solid lines represent fitted line with a one-term Fourier function. n>200 in each stage. (C) EGFP intensity in the entire mesenchyme for E12.5 and within the cluster from E13.5 onward. Welch’s two-sample t-test, E12.5-E13.5: p<0.001, E13.5-E14.5: p<0.001, E14.5-E15.5: p<0.001. n>100 in each stage. (D) Relative expression level of *Sox9, Col2a1*, and *Agc1* from E12.5 to E15.5. One-sample t-test, Sox9, E13.5: p=0.464, Col2a1, E13.5: p=0.171, Agc1, E13.5: p=0.158. Welch’s two-sample t-test, Sox9, E13.5-E14.5: p<0.001, E14.5-15.5: p=0.344. Col2a1, E13.5-E14.5: p=0.011, E14.5-15.5: p=0.048. Agc1, E13.5-E14.5: p=0.063, E14-E15: p=0.035. n=3.

We also quantified the expression levels of various cartilage matrix genes (the major determinants of chondrogenesis), including *Col2a1* and *Agc1*, as well as Sox9, by RT-qPCR analysis. As anticipated, *Sox9* expression showed a non-monotonic profile with a peak at E14.5, similar to that obtained from our imaging analyses of Sox9-EGFP (Fig. 2D). However, *Col2a1* and *Agc1* exhibited monotonic increases during progression through the developmental stages (Fig. 2D). These results suggest that other signaling inputs, together with Sox9, primarily regulate the expression of the cartilage matrix genes.

### Erk activity increases up to E14.5 and decreases thereafter, alongside Sox9 expression

Next, to examine the contribution of the Fgf-Erk axis, we quantified the spatiotemporal Erk activity in the developing trachea. For this purpose, we used a reporter mouse line that expresses a Förster resonance energy transfer (FRET)-based biosensor for Erk activity, which is localized in the nucleus (Harvey et al., 2008; Komatsu et al., 2011, 2018). Despite being designed for ubiquitous expression, the fluorescence signal of non-chondrogenic nodules was so dim that Erk activity could be quantified only in the chondrogenic nodules.

FRET imaging by two-photon microscopy revealed that Erk activity in the mesenchymal condensations increased 1.05-fold from E12.5 to E13.5, and 1.26-fold from E13.5 to E14.5, but decreased 0.89-fold from E14.5 to E15.5, (Fig. 3A and 3B). It is worth noting that this activity profile is similar to the expression profile of Sox9 (Fig. 2D). Treatment with PD0325901, an inhibitor for Mek (the kinase upstream of Erk), led to a significant decrease in Erk activity at all stages (Fig. 3B), suggesting that the level of Erk activity has a potential role in the developmental process.

**Figure 3.**
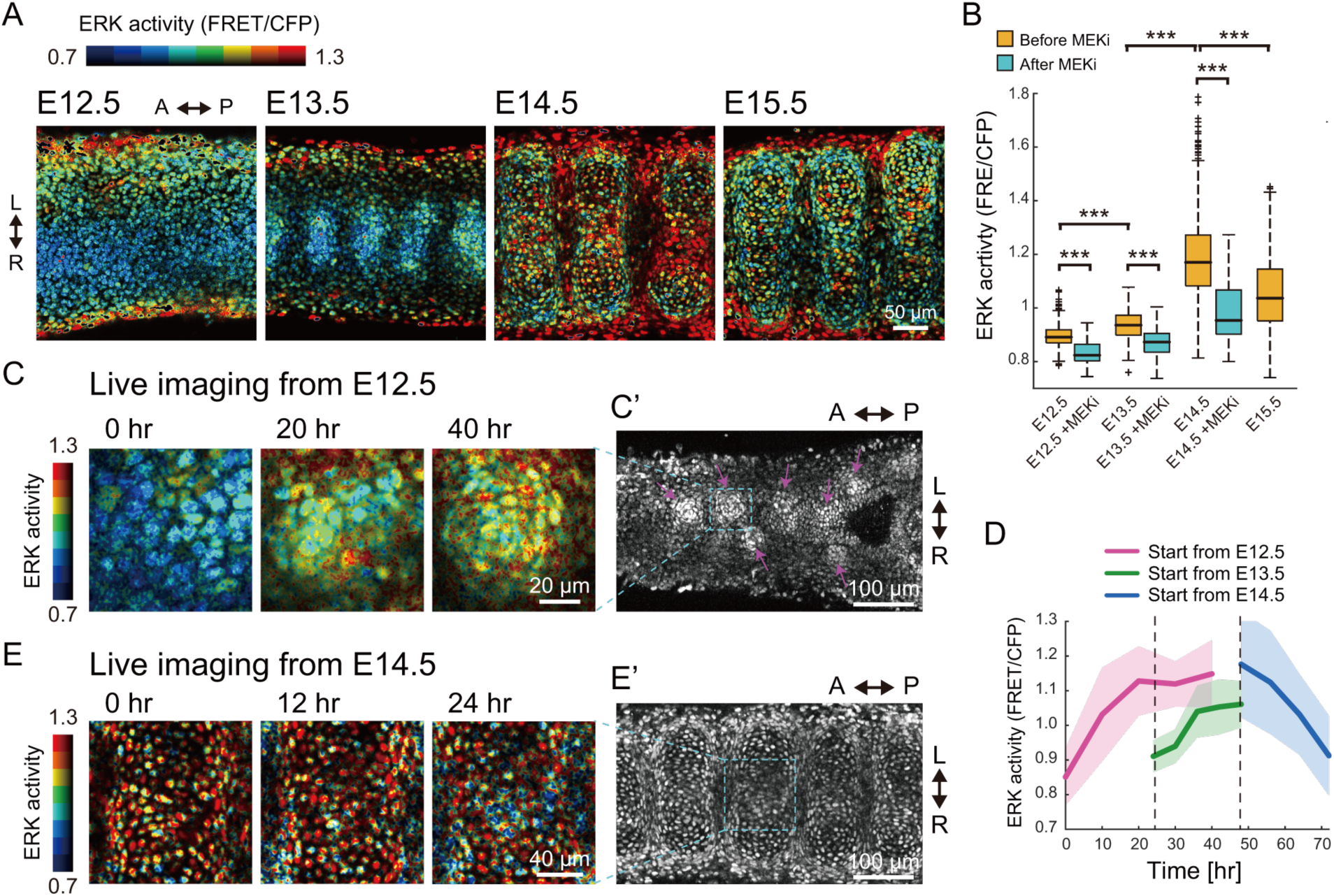
Erk activity profile in the developing trachea. (A) Erk activity map in the trachea on the frontal plane of the ventral side of the epithelium. Color denotes levels of Erk activity. Scale bar, 50 µm. L: left, R: right, A: anterior, P: posterior. (B) Erk activity in the condensed region before and after 120 min of treatment with the Mek inhibitor (500 nM). Welch’s two-sample t-test, p<0.001 in all groups. n>480 for each category from 3 different independent experiments. (C and C’) Time-lapse snapshots of Erk activity in the mesenchyme of an explant trachea cultured from E12.5. The far right panel in (C) corresponds to the dotted window in (C’). Magenta arrows indicate the mesenchymal condensations. Scale bars, 20 µm for (C) and 100 µm for (C’). (D) Time-series of Erk activity in nodule mesenchyme. Color denotes the different starting stages of the explant cultures. Explant tracheae were cultured from E12.5. Mean and s.d. n>70 from 2 independent experiments. (E and E’) Time-lapse snapshots of Erk activity in the mesenchyme of an explant trachea cultured from E14.5. The far right panel in (E) corresponds to the dotted window in (E’). Scale bars, 40 µm for (E) and 100 µm for (E’).

We also performed time-lapse imaging of the dissected tracheae using the FRET biosensor-expressing mice to continuously monitor the dynamics of the mesenchymal cells and the changes in Erk activity. In tracheae cultured from E12.5 embryos, the mesenchymal cells in the ventral epithelium formed some condensations and Erk activity gradually increased during this process (Fig. 3C and 3C’). Single cell quantification indicated that Erk activity increased to a maximum of 1.31-fold when cultured from E12.5 for 40 hours (Fig. 3D). The increase in Erk activity was also observed when E13.5 tracheae were cultured for 24 hours (1.17-fold, Fig. 3D). In contrast, Erk activity in the chondrogenic nodules decreased 0.78-fold when cultured from E14.5 (Fig. 3D, 3E, and 3E’). These observations confirmed that the gradual activation of Erk switches at E14.5 to inactivation.

### Erk activation is dispensable for mesenchymal condensation

We then explored the role of Erk activation in the development of tracheal cartilage. For these experiments, we treated tracheae dissected from E12.5 embryos with a Mek inhibitor under ex vivo culture conditions for one day. Since <u>how the nuclei change shape</u> in a way that is characteristic of mesenchymal condensation in the initial differentiation step occurring between E12.5 and E13.5 (Fig. 1A and 1B), we focused on assessing the shape and distribution of the mesenchymal nuclei, as well as Sox9 levels in the condensations. We observed elongated mesenchymal nuclei, that were distributed concentrically in tracheae treated with the Mek inhibitor, similar to the control samples (Fig. 4A, 4B, and 4D) and to the E13.5 samples (Fig. 1). Furthermore, the increase in Sox9 levels during mesenchymal condensation was confirmed in PD0325901-treated tracheae as well as in the control explants (Fig. 4A and 4B). These results indicate that the activation of Erk between E12.5 and E13.5 is dispensable for mesenchymal condensation, which is consistent with a previous report (Oh et al., 2000).

**Figure 4.**
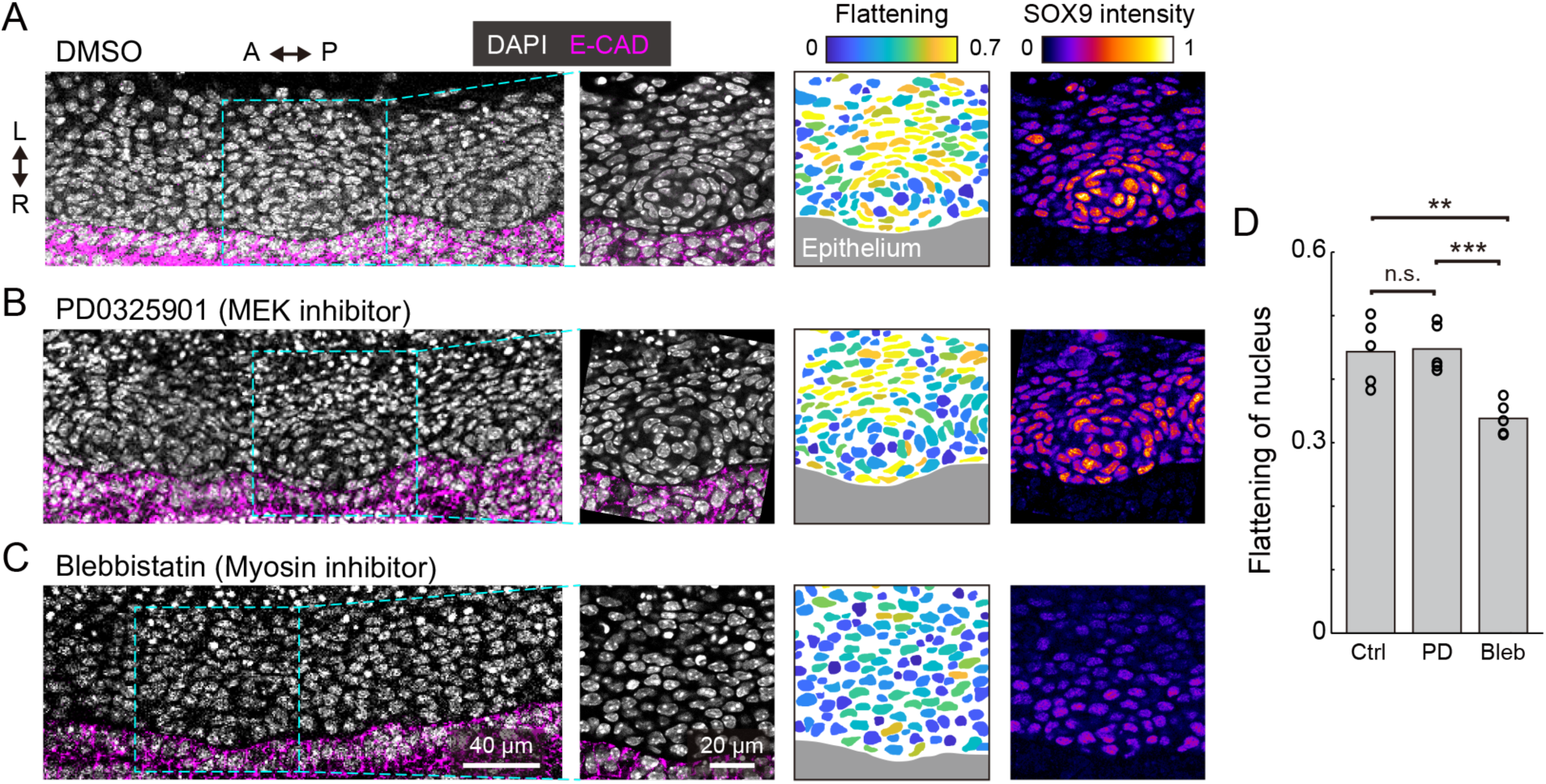
Erk activation in mesenchymal condensation. (A-C) Immunohistochemistry images of anti-E-cadherin (magenta) and anti-SOX9 (‘fire’ pseudocolor) with DAPI nuclear counterstaining (white) at the frontal section of the trachea, cultured ex vivo from E12.5 for 1 day with drug treatment (A: DMSO, B: PD0325901 at 500 nM, C: Blebbistatin at 30 µM). (1st and 2nd columns) Magnified images from the dotted windows in the 1st column show concentric mesenchymal condensation in the DMSO control and PD0325901-treated (A and B) but not in the blebbistatin-treated (C) tracheae. (3rd and 4th columns) Color denotes the flattening of nuclei and Sox9 intensity, respectively, corresponding to the images from the 2nd column. Marked increases in Sox9 levels were observed in the DMSO control and PD0325901-treated (A and B) but not in the blebbistatin-treated (C) tracheae. Scale bars, 40 µm (1st column) and 20 µM (2nd-4th column). L: left, R: right, A: anterior, P: posterior. (D) Flattening of nuclei in response to different treatments. Each point represents the mean value within a sample. Welch’s two-sample t-test, Ctrl-PD: p=0.885. Ctrl-Bleb: p=0.007, PD-Bleb: p=0.001. n=5.

We also examined the effect of mesenchymal condensation on the suppression of cell-generated contractile forces, since it has been shown that cell contraction drives mesenchymal condensation in the development of other organs (Mammoto et al., 2011; Shyer et al., 2017). Tracheae dissected at E12.5 were treated with blebbistatin, an inhibitor of non-muscle myosin II, and cultured for one day. Most mesenchymal nuclei remained round, and the tissue did not exhibit clear condensations or increased Sox9 expression near the epithelium (Fig. 4C and 4D), suggesting that cell-generated contractile forces are required for mesenchymal condensations in tracheal development, but that Erk activation is dispensable for the regulation of contractile forces in this process.

### Erk inactivation promotes expression of cartilage matrix genes

To investigate the role of Erk activation in tracheal cartilage formation from E13.5 onwards, we measured the expression levels of *Sox9* and several cartilage matrix genes, which show a marked elevation in expression at this stage, in response to treatment with the Mek inhibitor (Fig. 2D). The tracheae were dissected at E13.5 and cultured for 2 days in the presence of PD0325901 under ex vivo conditions. The expression of *Spry2*, an indicator of Ras–Erk cascade activity (Mason et al., 2006), was decreased by PD0325901 treatment, confirming the inhibition of Erk activation in this assay (Fig. 5A). The expression of *Sox9* was unaffected, as anticipated from the data shown in Fig. 4. We also confirmed that Erk inactivation did not affect *Sox9* expression using imaging measurements of Sox9-EGFP tracheae (Fig. S2). These data regarding Erk signaling are similar to those from a previous study which reported that inactivation of *Fgf10* did not affect *Sox9* expression (Sala et al., 2011). Interestingly, Erk inhibition significantly upregulated the expression of *Col2a1* and *Agc1* (Fig. 5A). These findings suggest that Erk inactivation promotes gene expression of these cartilage matrix proteins in a manner independent of *Sox9* expression.

**Figure 5.**
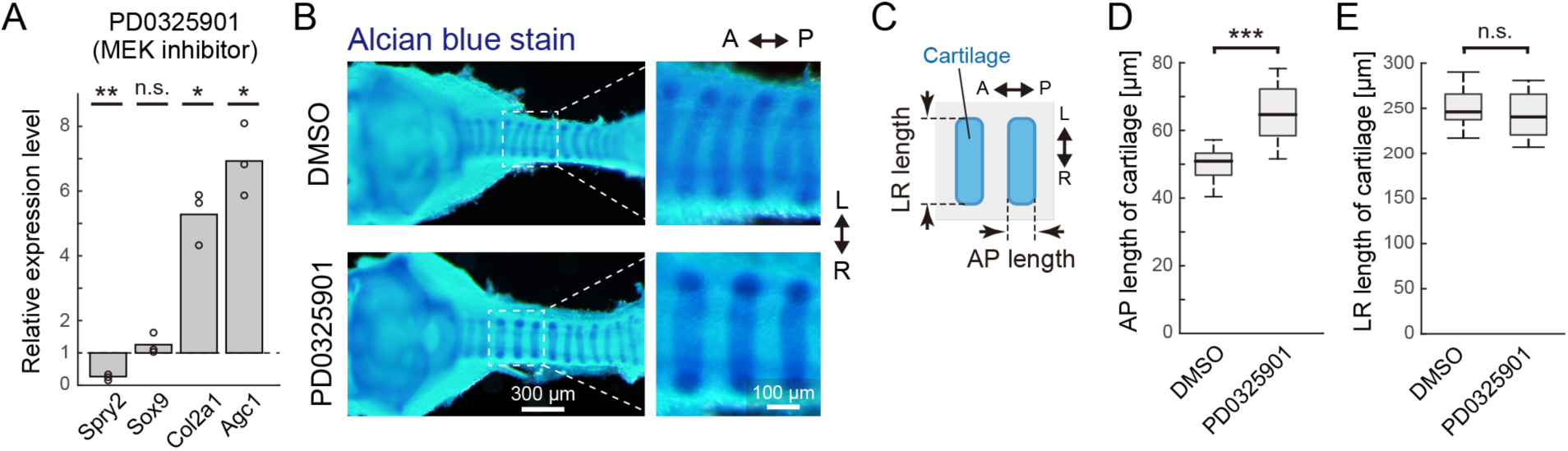
Erk inactivation in cartilage matrix gene expression. (A) Relative gene expression in tracheae cultured ex vivo for 2 days from E13.5 with the PD0325901, compared to DMSO control, obtained by qPCR. Welch’s one-sample t-test. *Spry2*: p=0.006, *Sox9*: p=0.255, *Col2a1*: p=0.013, *Agc1*: p=0.012. n=3. (B) Morphological change in tracheal cartilage treated with PD0325901. Cartilage rings were stained with Alcian blue and photographed on a dissecting microscope. Scale bars, 200 µm (left) and 100 µm (right). (C) Schematics showing the antero-posterior (AP) length and left-right (LR) length of the tracheal cartilage rings. The 3rd to 6th tracheal rings below the cricoid ring were measured. (D) AP length of cartilage rings exposed to the DMSO control or PD0325901. Welch’s two-sample t-test. p<0.001. n=18. (E) LR length of cartilage rings exposed to the DMSO control or PD0325901. Welch’s two-sample t-test. p=0.281. n=18. L: left, R: right, A: anterior, P: posterior.

We next examined the impact of Erk inactivation on the tracheal cartilage phenotype. To this end, PD0325901 was administered to pregnant mice from E13.5 periodically for two days by oral gavage. The morphology of tracheal cartilage rings from dissected embryos was assessed by Alcian blue staining (Fig. 5B), and the tracheae were evaluated in terms of antero-posterior (AP) and left-right (LR) lengths (Fig. 5C). Erk inactivation significantly increased the AP length (Fig. 5D), while it did not affect the LR length (Fig. 5E), indicating that cartilage matrix accumulation was enhanced due to Erk inactivation. Together, our results suggest that active Erk represses the transcriptional activity of the cartilage matrix genes, thereby influencing the AP length of the cartilage rings.

### Incoherent feedforward loop can explain the regulation of Sox9-Erk-cartilage matrix genes

Our results so far present two pathways which have antagonistic effects on the transcriptional regulation of the cartilage matrix genes, namely activation via Sox9 and repression via Erk. Furthermore, since Sox9 expression and Erk activity exhibited similar non-monotonic temporal profiles (Fig. 2D and 3B), it suggests that there may be a common upstream regulator X (Fig. 6A, left) or positive regulation of Erk by Sox9 (Fig. 6A, right). These network topologies are examples of a so-called incoherent feedforward loop (IFFL), known as a network motif, which represent recurrent patterns in transcriptional regulations (Shen-Orr et al., 2002; Mangan and Alon, 2003). We therefore considered the question of how these regulatory networks might function in tracheal development.

**Figure 6.**
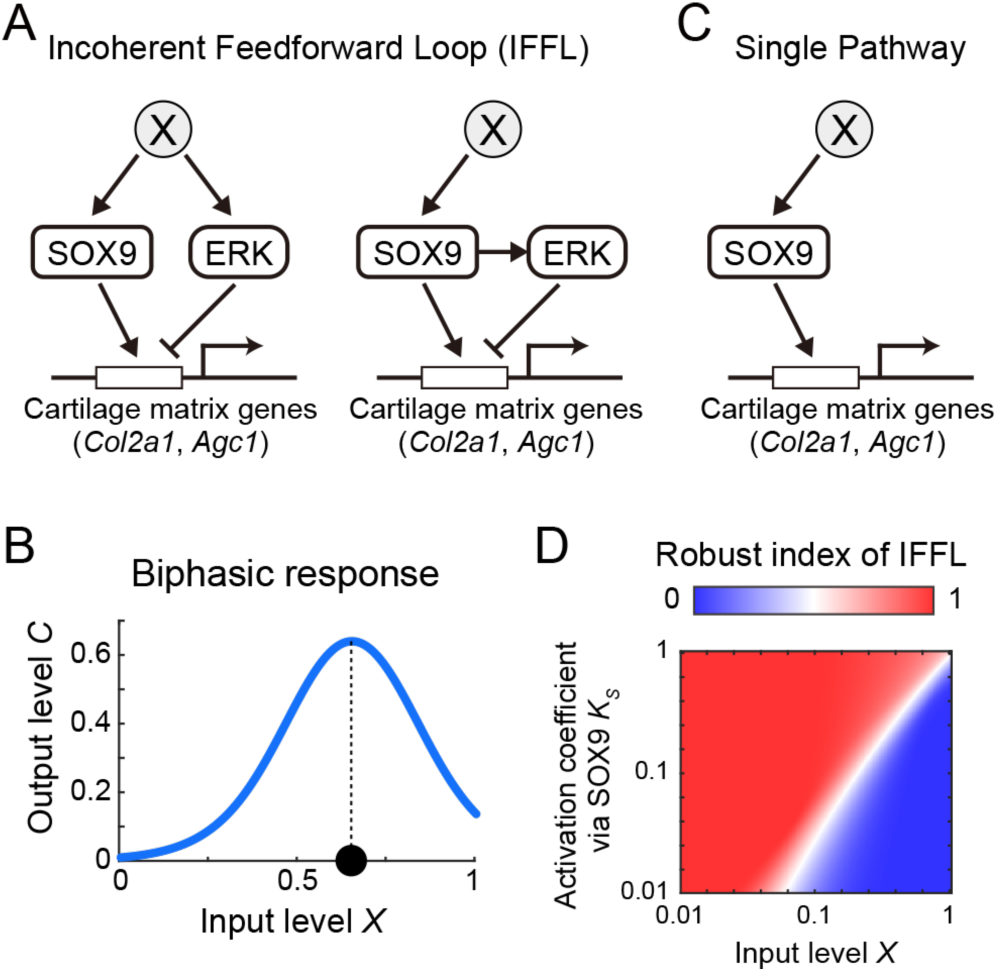
Model of incoherent feedforward loop exploring the regulation of cartilage matrix genes by Sox9/Erk. (A) Possible network for cartilage matrix gene expression. (B) Input-output response of IFFL showing biphasic response. (C) Hypothetical single pathway. (D) Robust index of IFFL with regards to *X* and 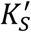.

We modeled the possible core regulatory networks (Fig. 6A) with a linear relation between X, Sox9, and Erk (see (2) Mathematical analysis in Materials and Methods) because of the similar temporal variations, and found that, in either case, the expression level of the cartilage matrix genes in the steady state *Ĉ* converged to the following form:

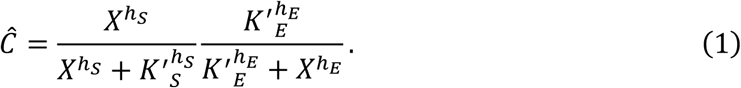

The first term represents the activation by Sox9 with the coefficient 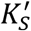, and the second term represents the repression via Erk with the coefficient 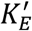, each of which is modeled by the Hill function with coefficient *h*_*S*_ or *h*_*E*_, respectively, with input level *X* as the simplest form. The function of Eq. 1 shows a biphasic inverse U-shape response, in which the output level *Ĉ* increases to a high level, and then decreases, as input level *X* increases, meaning that the output level reaches at its maximum at an intermediate input level (Fig. 6B and Fig. S3). From Eq. 1, the biphasic response modeled by the IFFL can explain the mapping of the non-monotonic temporal profiles of both Sox9 and Erk to the monotonic increases of the cartilage matrix genes (Fig. 2E).

We next considered the role of Erk as a negative regulator of the pathways involved in chondrogenesis and queried the benefit of an IFFL involving Erk as a core regulatory design, compared with a single pathway regulated only by Sox9. To answer these questions, we analyzed the robustness of the steady state level of cartilage matrix gene expression *Ĉ* with respect to the input level *X* in the IFFL compared to the hypothetical single pathway (Fig. 6A and 6C). To do this, we calculated the parameter sensitivity coefficient (Goldbeter and Koshland, 1981) of *Ĉ* with respect to *X* in both the IFFL (*S*_1_) and in the single pathway (*S*_0_), and obtained a condition of parameters which satisfies *S*_1_> *S*_0_. That is, IFFL is more robust than the single pathway for input variations in the following inequality:

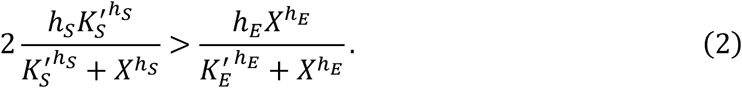

Here we introduced a fraction of parameter sets that satisfy Eq. 2, designated Robust index of IFFL. When this value is more than 0.5, the IFFL is a more robust design compared with the single pathway, and vice versa. We then numerically investigated the dependency of the parameters *X* and 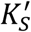, both of which are common to the IFFL and the single pathway, without fixing other parameters, and found that the parameter space where the index is more than 0.5 was larger than the area representing an index below 0.5 (Fig. 6D). Collectively, our model analyses show that adopting Erk as a negative regulator of the regulatory chondrogenic pathways makes the system more robust against variations of the upstream regulator of Sox9 during chondrogenesis.

## Discussion

In this study, we demonstrate that there is an inverted U-shaped temporal change in Sox9 expression and Erk activity, while the expression of target cartilage matrix genes monotonically increases, during murine tracheal development. Using assays with a Mek inhibitor, we found that inhibiting Erk activity significantly promotes the expression of cartilage matrix genes. The role played by Erk during cartilage differentiation has been controversial in several *in vitro* studies (Murakami et al., 2000; Oh et al., 2000; Bobick and Kulyk, 2004). Our results show that the Erk signaling pathway functions as an inhibitory regulator of tracheal cartilage differentiation, which has also been demonstrated in embryonic chicken limb buds (Oh et al., 2000; Bobick and Kulyk, 2004).

Our findings led us to propose a model in which cartilage differentiation is controlled by an IFFL via positive and negative contributions of Sox9 and Erk activity, respectively. To delineate the core regulatory pathways, we tested two networks, one of which included a common upstream regulator without interactions between Sox9 and Erk, while the other network included a positive regulation of Erk by Sox9 (Fig. 6A), although both of them resulted in the same class of output in the steady state (Eq. 1). It has previously been demonstrated that the expression level of *Fgf10* does not change significantly even in *Sox9* knockout mice (Turcatel et al., 2013). Furthermore, there is evidence to imply that Shh, an upstream regulator of *Sox9*, could be a potential activator of Fgf10 in tracheal development (Sala et al., 2011), and it raises the possibility that Shh could also be the common upstream regulator of Sox9 and Erk. It is therefore likely that the former regulatory network is adopted during tracheal ring formation.

In this study, we primarily focused on the expression of the cartilage matrix genes as an output, but the spatiotemporal dynamics of Sox9 expression and Erk activity are likely to play roles in other aspects of cartilage differentiation. From E14.5 to E15.5, we observed a decrease in *Sox9* expression (Fig. 2C). This may reflect the fact that the downregulation of SOX9 has multiple roles in the progression of cartilage differentiation, such as the transition of proliferating chondrocytes to a hypertrophic state, and cartilage vascularization (Akiyama et al., 2002; Hattori et al., 2010; Lefebvre et al., 2019). We also observed that endogenous Erk activation was maintained at a relatively low level from E12.5 to E13.5 (Fig. 3B) and did not affect mesenchymal condensation, an initial step in tracheal ring patterning (Fig. 4B). Since *Fgf10* overexpression in the mesenchyme between E11.5 and E13.5 was shown to cause disruption of the tracheal ring patterns (Sala et al., 2011), it is possible that Erk activity at below-moderate levels may be a key factor in normal tracheal development. Future studies should clarify the physiological importance of temporal Erk activity dynamics in this process. In addition, further systematic investigations of other signaling pathways, such as Wnt and bone morphogenetic protein (Bmp) signaling (Snowball et al., 2015; Kishimoto et al., 2018), as well as physical interactions with epithelia and smooth muscles (Hines et al., 2013; Yin et al., 2018), would improve our understanding of tracheal development and in vitro organoid systems (Kishimoto et al., 2019; Conway et al., 2020).

## Acknowledgements

This work was supported by the JSPS KAKENHI 17KT0107 and 19H00993, 16H06280, by the JST PRESTO JPMJPR1949, by the CREST JPMJCR1654, by the AMED 19gm5010003h0003, and by the Medical Research Support Center of Kyoto University. We would like to thank Yu Kurata and Akane Kusumi for technical assistances, and Keisho Hirota and Mitsuru Morimoto for fruitful discussion.

## Author Contributions

Conceptualization, T.Y., T.H.; Methodology, T.Y., T.H.; Software, T.H.; Validation, T.Y., T.H.; Formal analysis, T.Y., T.H.; Investigation, T.Y., T.H.; Resources, M.M., T.H., T.H.; Data curation, T.Y., T.H.; Writing - original draft, T.Y., T.H.; Writing - review & editing, M.M., T.H.; Visualization, T.H.; Supervision, M.M., T.H.; Project administration, T.H.; Funding acquisition, M.M., T.H.

## Materials and Methods

### (1) Experiments and quantification

#### Animals

For FRET imaging, we used transgenic mice that ubiquitously express an ERK biosensor with a long flexible linker, which has been described elsewhere (Harvey et al., 2008; Komatsu et al., 2011, 2018). Sox9-EGFP mice were provided from RIKEN BRC through the National BioResource Project of the MEXT/AMED, Japan (RBRC05651). Otherwise, we used ICR mice purchased from Japan SLC, Inc. The midnight preceding observation of a plug was designated as embryonic day 0.0 (E0.0), and all mice were sacrificed by cervical dislocation to minimize suffering. All the animal experiments were approved by the local ethical committee for animal experimentation (MedKyo 19090 and 20081) and were performed in compliance with the guide for the care and use of laboratory animals at Kyoto University.

#### Antibodies

The following primary and secondary antibodies were used for immunofluorescence: anti-E-cadherin rat antibody (#3195, 1:100 dilution, Cell Signaling Technology), anti-SOX9 rabbit antibody (#AB5535-25UG, 1:200 dilution, Merck Millipore), Alexa Fluor 546-conjugated goat anti-rat IgG (H+L) antibody (#A11081, 1:1000 dilution), Alexa Fluor 647-conjugated goat anti-rat IgG (H+L) antibody (#A21247, 1:1000 dilution), Alexa Fluor 647-conjugated goat anti-rabbit IgG (H+L) antibody (#A32733, 1:1000 dilution)(all Thermo Fisher Scientific).

#### Whole-tissue fluorescence staining and imaging

Staining and optical clearing of dissected tracheae were performed as described in a previous study (Hirashima and Adachi, 2015). Briefly, the samples were fixed with 4% PFA in PBS overnight at 4°C. For anti-SOX9 staining, the samples were incubated in 25 mg/mL hyaluronidase (Nacalai Tesque, #18240-36) for 1 h at 37°C, to digest hyaluronic acid. The samples were then blocked in 10% normal goat serum (Abcam, #ab156046) diluted in 0.1% Triton X-100/PBS (PBT) for 3 h at 37°C. The samples were treated with primary antibodies overnight at 4°C, washed in 0.1% PBT, and subsequently incubated in secondary antibodies conjugated to either Alexa Fluor 546 or Alexa Fluor 647 overnight at 4°C. DAPI was used for nuclear counterstaining (Dojindo Molecular Technologies, #D523-10, 1:200 dilution). The samples were mounted with 10 µL of 1% agarose gel onto a glass dish (Greiner Bio-One, #627871) for stable imaging. Then, the samples were immersed in CUBIC-R+ (Tokyo Chemical Industry Co., # T3741) solution for optical clearing. Images were acquired using the confocal laser scanning platform Leica TCS SP8 equipped with the hybrid detector Leica HyD, using a ×40 objective lens (NA = 1.3, WD = 240 μm, HC PL APO CS2, Leica) and the Olympus FluoView FV1000 with a ×30 objective lens (NA = 1.05, WD = 0.8 mm, UPLSAPO30XS, Olympus).

#### Alcian blue staining

The dissected tracheae were fixed in 4% PFA in PBS overnight at 4°C, and stained in Alcian Blue Solution (FUJIFILM Wako Pure Chemical Corporation, # 013-13801) for 60 min at 23°C. The samples were then washed with 20% acetic acid in PBS overnight at 23°C, and finally clarified in 50% glycerol in PBS for 2 hours at 37°C. The samples were visualized by the stereo microscopy (SZX16, Olympus).

#### Explant cultures

The dissected tracheae were mounted on a 35 mm glass dish (Greiner, #627871) with 30 μL of growth factor-reduced Matrigel (Corning, #356231), and filled with 500 µL of DMEM culture medium including FluoroBrite (Thermo Fischer Scientific, #A1896701) with 1% GlutaMAX (Thermo Fischer Scientific, #35050061). The samples were incubated at 37°C under 5% CO_2_.

#### Drug administration

For the drug administration under ex vivo culture conditions, (-)-Blebbistatin (FUJIFILM Wako Pure Chemical Corporation, #021-17041) and PD0325901 (FUJIFILM Wako Pure Chemical Corporation, #162-25291) were mixed in the culture medium. Equivalent amounts of DMSO were used as a vehicle control for each drug. For administration by oral gavage, PD0325901 (ChemieTek, #CT-PD03) in 30% PEG400, 0.5% Tween 80 in PBS was administered to pregnant mice at a dose of 25 mg/kg body weight twice a day (at 8:00am and 6:00pm) for two days from E13.5, i.e., 4 times in total.

#### Live imaging for explants

The samples were prepared for explant culture as described above and placed into an incubator-integrated multiphoton fluorescence microscope system (LCV-MPE, Olympus) with a ×25 water-immersion lens (NA=1.05, WD=2 mm, XLPLN25XWMP2, Olympus) and an inverted microscope (FV1200MPE-IX83, Olympus) with a ×30 silicone-immersion lens (NA=1.05, WD=0.8 mm, UPLSAPO30XS, Olympus). The excitation wavelengths were set to 840 nm or 930 nm, for the CFP of the Erk FRET biosensor and EGFP, respectively (InSight DeepSee, Spectra-Physics). The filter sets used were as follows; IR cut filter: RDM690 (Olympus), dichroic mirrors: DM505 and DM570 (Olympus), and emission filters: BA460-500 for CFP, BA520-560 for FRET, and BA495-540 for EGFP detection (Olympus).

#### Quantitative RT-PCR

Total RNA was extracted using the RNeasy Mini Kit (Qiagen, #74104), and cDNA was reverse transcribed using the High-Capacity cDNA Reverse Transcription Kit (Thermo Fisher Scientific, #4368814), according to the manufacturer’s instructions. qPCR was performed using the StepOne real-time PCR system (Applied Biosystems) with PowerUp SYBR Green Master Mix (Thermo Fisher Scientific, #A25742). Primer sequences were as follows: *Agc1* forward 5’-GGTCACTGTTACCGCCACTT-3’ and reverse 5’-CCCCTTCGATAGTCCTGTCA-3’; *Col2a1* forward 5’-CTACGGTGTCAGGGCCAG-3’ and reverse GCAAGATGAGGGCTTCCATA-3’; *Hprt1* forward 5’-TCAGTCAACGGGGGACATAAA-3’ and reverse GGGGCTGTACTGCTTAACCAG-3’; *Sox9* forward 5’-AGGAAGCTGGCAGACCAGTA-3’ and reverse TCCACGAAGGGTCTCTTCTC-3’; *Spry2* forward 5’-AGAGGATTCAAGGGAGAGGG-3’ and reverse 5’-CATCAGGTCTTGGCAGTGTG-3’. Relative expression levels were calculated using the ΔΔCT method with *Hprt1* expression as the internal control.

#### FRET image analysis

The median filter of a 3×3 window was processed to remove shot noises, and the background signal was subtracted each in FRET and CFP channel. Then, the ratio of FRET intensity to the CFP intensity was calculated using a custom-designed MATLAB (MathWorks) script.

#### Quantification of nuclear shape

The DAPI staining images were smoothened using the median and gaussian filters, and the nuclei were then manually extracted. Flattening or ellipticity is defined as 1-*b*/*a*, where *a* and *b* are the major and the minor axis length of the best fitting ellipse, respectively. The value is 0 for a circle, and it approaches 1 as it is compressed. All processing was done using ImageJ.

#### Statistical hypothesis testing

Statistical tests, sample sizes, test statistics, and *P*-values are described in the main text. *P*-values of less than 0.05 were considered to be statistically significant in two-tailed tests, and were classified into 4 categories; * (*P<*0.05), ** (*P*<0.01), *** (*P*<0.001), and n.s. (not significant, i.e., *P* ≥ 0.05).

#### Software

For digital image processing, MATLAB (MathWorks) and ImageJ (National Institute of Health) were used. For graphics, MATLAB (MathWorks), Imaris (Bitplane) and ImageJ (National Institute of Health) were used. MATLAB (MathWorks) was used for statistical analysis and Mathematica (Wolfram Research) for mathematical analysis.

#### Graph

For the boxplot, the central mark indicates the median, and the bottom and top edges of the box indicate the 25th and 75th percentiles, respectively. The whiskers extend to the most extreme data points not considered outliers, and the outliers are plotted individually using the ‘+’ symbol. All of the graphs were prepared in MATLAB.

### (2) Mathematical analysis Modeling

For model construction, we let *X, S, E, C* be input level, Sox9 concentration, Erk activity, and the expression level of cartilage matrix genes, respectively. Since Sox9 expression and Erk activity showed similar temporal profiles, they would be regulated linearly by input level *X*. Thus,

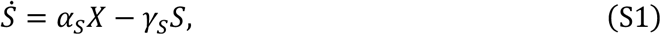

and

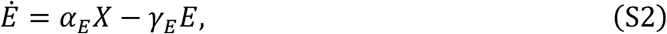

where *α* and *γ* denote production rate and decay rate, respectively. Regarding the alternative regulation, i.e., X indirectly, but not directly, regulates Erk activity through Sox9, the regulation of Erk activity can be represented, instead of Eq. S2, as

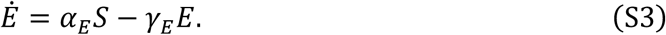

As Sox9 and active Erk antagonistically regulate the expression level of cartilage matrix genes, the dynamics of *C* can be represented in the simplest form as follows:

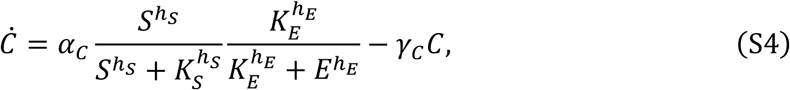

where *K*_*S*_ is the activation coefficient via Sox9, *K*_*E*_ is the repression coefficient via Erk, and *h* denotes the Hill coefficient. Both combinations of Eqs. S1, S2, and S4, and Eqs. S1, S3, and S4 led to the following function of *C* in the steady state:

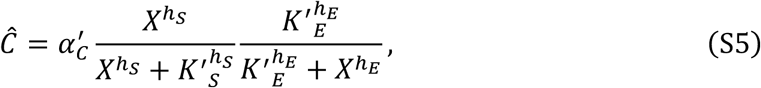

where parameters with prime represent integrated parameters, and the case with 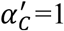 is shown as the Eq. 1 in the main text.

#### Parameter ranges in numerical investigation

Owing to the importance of relativity for *X*, 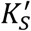, and 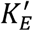, we set those ranges as 0.01 to 1.0. For *h*_*S*_ and *h*_*E*_, we set the range from 1 to 5 to consider non-linearity of reactions. 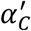 was arbitrarily set to 1.

#### Parameter dependence in the steady state

Assuming *h*_*S*_ = *h*_*E*_ = 1 for simplicity and feasibility of analysis, we found that the input value at the output peak *X*^*^ was determined by the two parameters 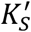 and 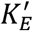as follows

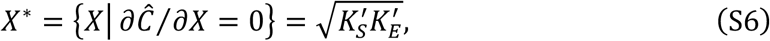

which corresponded to the numerical results (Fig. S3B’). Also, the function of *Ĉ* was convex upward at the peak as

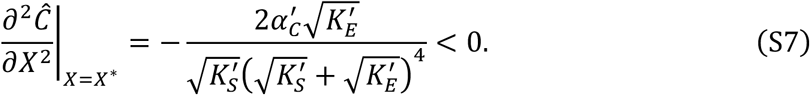

Moreover, the maximum level of *Ĉ* is represented as

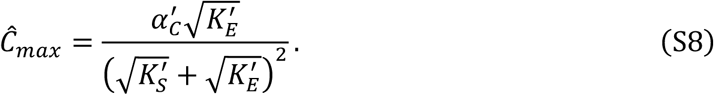

This clearly indicates that the output peak level decreases with increases in 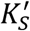 and it increases with increasing 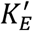, corresponding to the numerical results (Fig. S3B’).

#### Sensitivity analysis for robustness

The parameter sensitivity coefficient with respect to the input value *X*, denoted *S*(*Ĉ, X*), is defined as the relative change in *Ĉ* for a given small relative change in *X*:

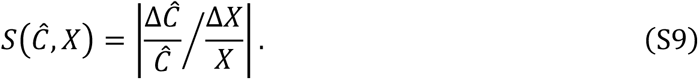

From Eqs. S5 and S9, we obtained the explicit form *S* in the IFFL regulation.

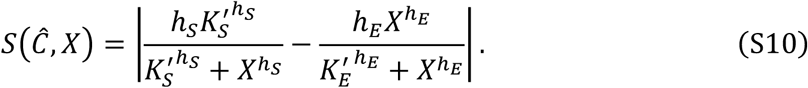

As an alternative regulation, we assumed a hypothetical single pathway regulation, instead of Eq. S4,

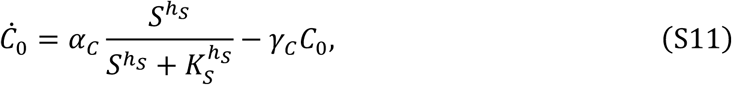

and from Eqs. S9 and S11, we obtained the parameter sensitivity coefficient for a single pathway as follows:

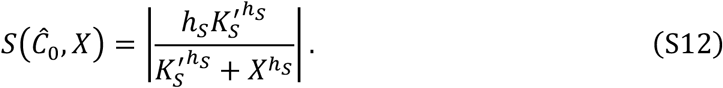

We then evaluated the robustness of the IFFL compared to the single pathway by the inequality of the sensitivity coefficients:

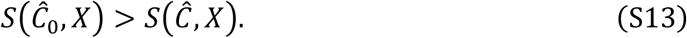

With Eqs. S10 and S12, and S13, Eq. S13 reached the inequality condition shown in Eq. 2 in the main text.

## Supplemental figure legends

**Figure S1.**
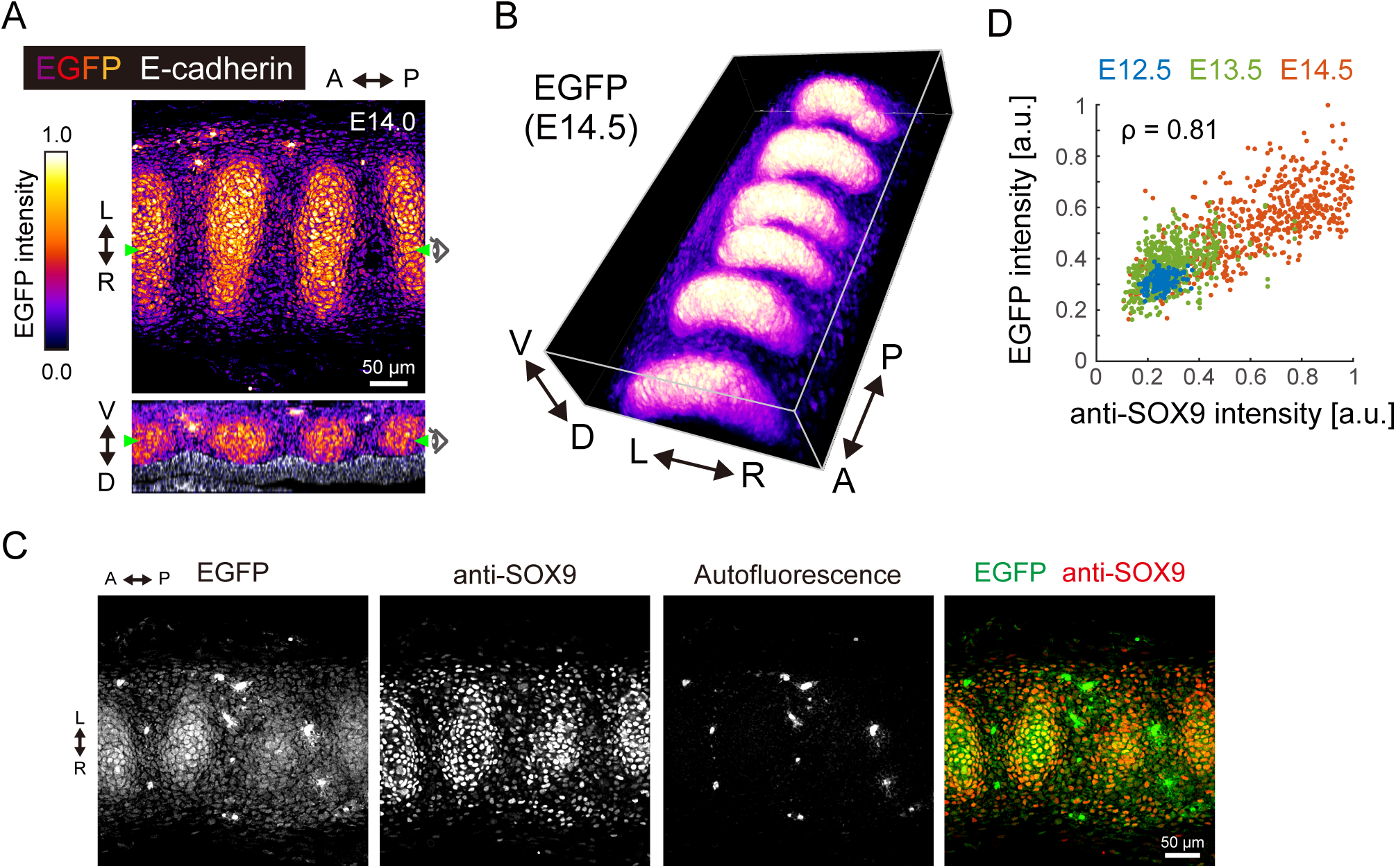
Sox9-EGFP profile in the developing murine trachea. (A) A 3D-rendered image of Sox9-EGFP at E14.5, showing the C-shaped tracheal rings on the ventral side. L: left, R: right, A: anterior, P: posterior, V: ventral, D: dorsal. (B) Simultaneous visualization of EGFP (‘fire’ pseudocolor) and anti-E-cadherin (white) at E14.0. High EGFP expression cell clusters located between the ridges of wavy tracheal epithelium. Scale bar, 50 µm. (C) Simultaneous visualization of EGFP (green) and anti-SOX9 (red) at E14.5. Scale bar, 50 µm. (D) Relationship between EGFP and anti-SOX9 intensity from E12.5 to E14.5. Pearson’s linear correlation coefficient: ρ=0.81.

**Figure S2.**
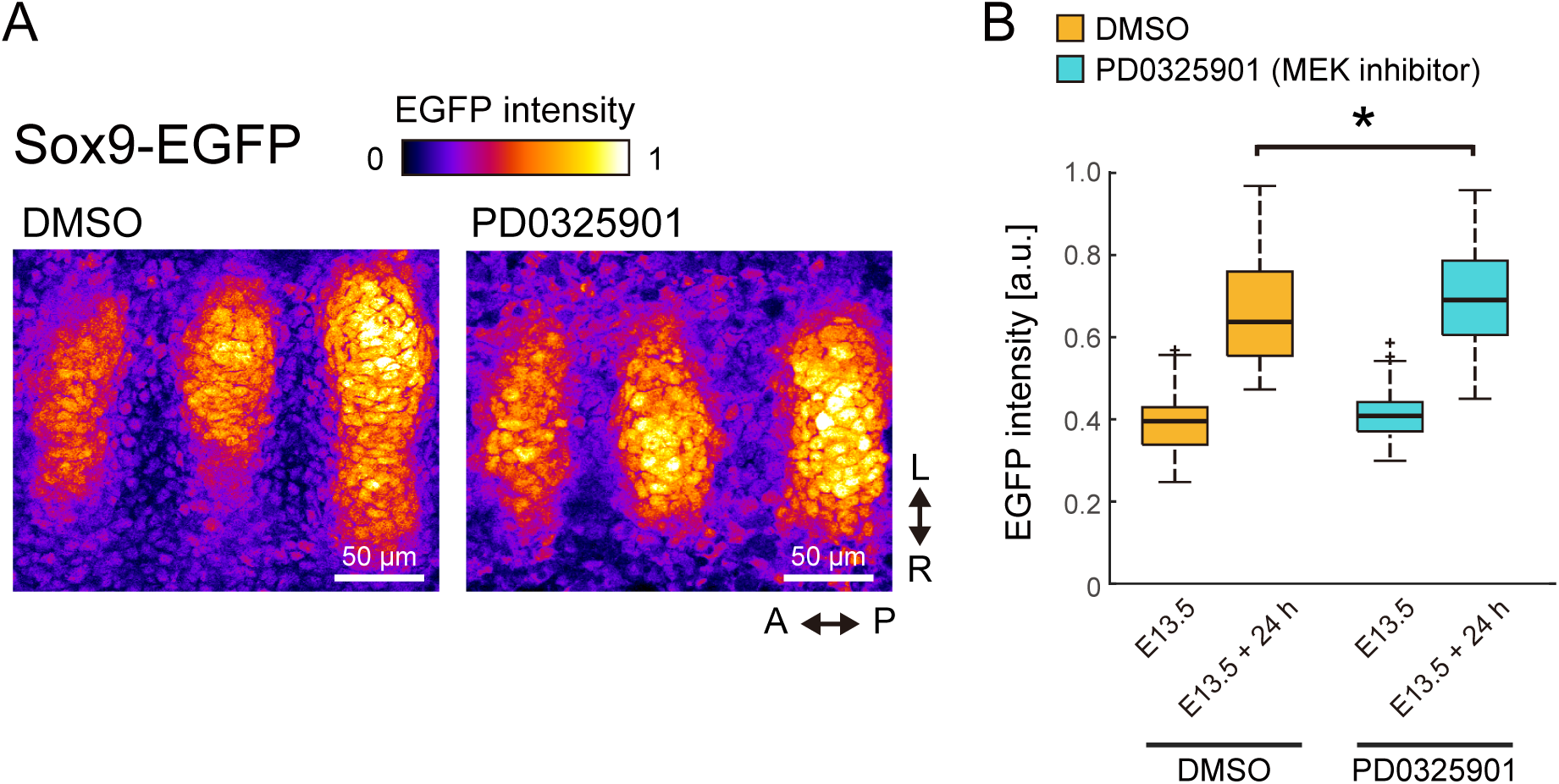
Effect of Erk inactivation on Sox9 expression. (A) Tracheae dissected at E12.5 from Sox9-EGFP mice were cultured for 1 day, together with PD0325901 (500 nM). EGFP intensity was measured by two-photon microscopy. (B) EGFP intensity with DMSO and PD0325901. Welch’s two-sample t-test, p=0.012. n=60.

**Figure S3.**
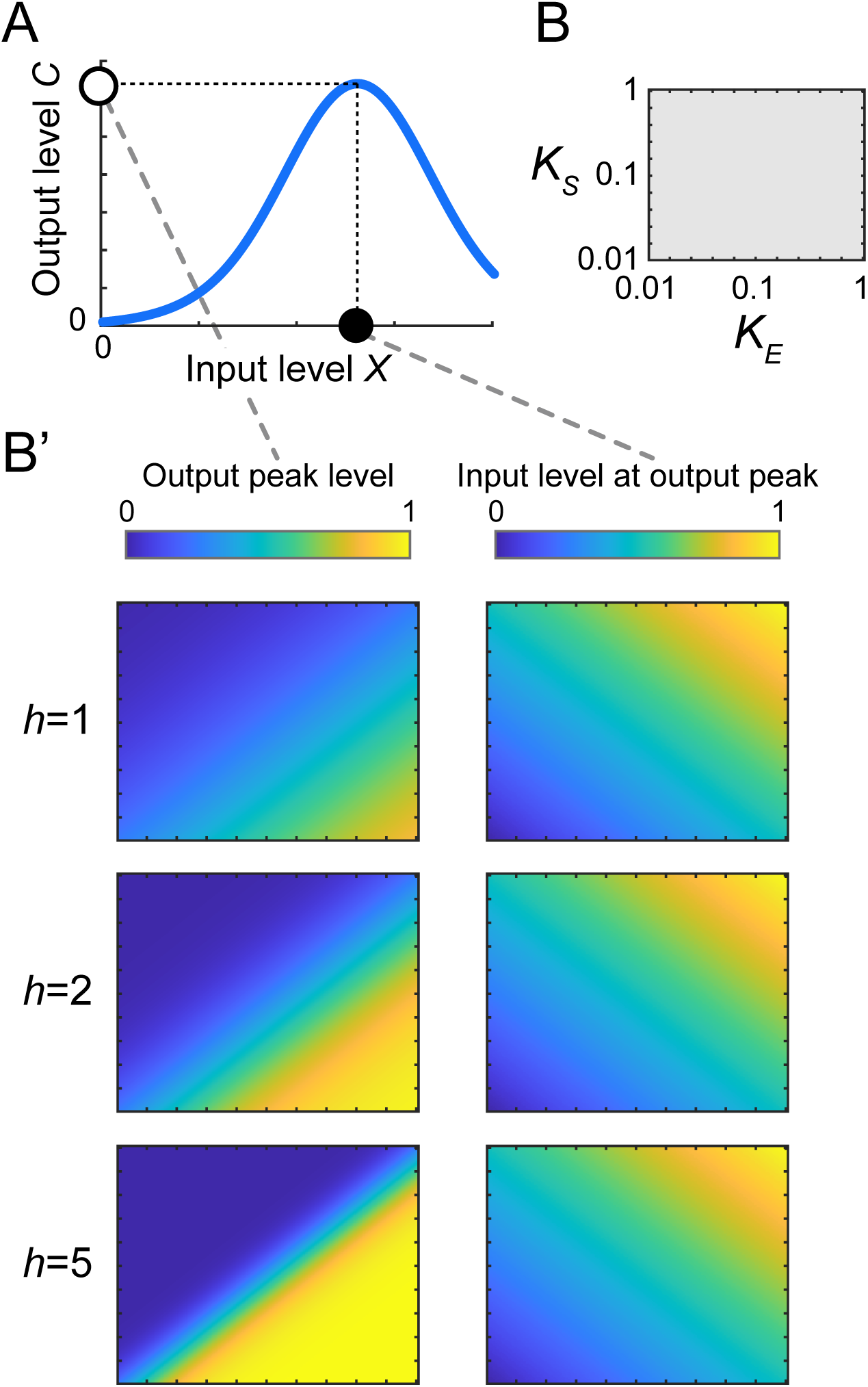
Parameter dependence of the model. (A) Schematics showing the input-output relationship. (B and B’) Parameter dependence of 1) the output peak level and 2) the input level at the output peak. Axes are shown in B.

## References

Akiyama, H., Chaboissier, M. C., Martin, J. F., Schedl, A., and De Crombrugghe, B. (2002). The transcription factor Sox9 has essential roles in successive steps of the chondrocyte differentiation pathway and is required for expression of Sox5 and Sox6. Genes Dev. doi: 10.1101/gad.1017802.

Arooj Sher, Z., and J Liu, K. (2016). Congenital tracheal defects: embryonic development and animal models. AIMS Genet. doi: 10.3934/genet.2016.1.60.

Bi, W., Deng, J. M., Zhang, Z., Behringer, R. R., and De Crombrugghe, B. (1999). Sox9 is required for cartilage formation. Nat. Genet. doi: 10.1038/8792.

Bi, W., Huang, W., Whitworth, D. J., Deng, J. M., Zhang, Z., Behringer, R. R., et al. (2001). Haploinsufficiency of Sox9 results in defective cartilage primordia and premature skeletal mineralization. Proc. Natl. Acad. Sci. U. S. A. doi: 10.1073/pnas.111092198.

Bobick, B. E., and Kulyk, W. M. (2004). The MEK-ERK Signaling Pathway Is a Negative Regulator of Cartilage-specific Gene Expression in Embryonic Limb Mesenchyme. J. Biol. Chem. doi: 10.1074/jbc.M309805200.

Boucherat, O., Nadeau, V., Charron, J., Berube-Simard, F.-A., and Jeannotte, L. (2015). Crucial requirement of ERK/MAPK signaling in respiratory tract development. Development. doi: 10.1242/dev.131821.

Cardoso, W. V, and Lü, J. (2006). Regulation of early lung morphogenesis: questions, facts and controversies. Development 133, 1611–1624. doi: 10.1242/dev.02310.

Conway, R. F., Frum, T., Conchola, A. S., and Spence, J. R. (2020). Understanding Human Lung Development through In Vitro Model Systems. BioEssays 42, 2000006. doi: 10.1002/bies.202000006.

Goldbeter, A., and Koshland, D. E. (1981). An amplified sensitivity arising from covalent modification in biological systems. Proc. Natl. Acad. Sci. U. S. A. doi: 10.1073/pnas.78.11.6840.

Goldring, M. B., Tsuchimochi, K., and Ijiri, K. (2006). The control of chondrogenesis. J. Cell. Biochem. doi: 10.1002/jcb.20652.

Han, Y., and Lefebvre, V. (2008). L-Sox5 and Sox6 Drive Expression of the Aggrecan Gene in Cartilage by Securing Binding of Sox9 to a Far-Upstream Enhancer. Mol. Cell. Biol. doi: 10.1128/mcb.00695-08.

Harvey, C. D., Ehrhardt, A. G., Cellurale, C., Zhong, H., Yasuda, R., Davis, R. J., et al. (2008). A genetically encoded fluorescent sensor of ERK activity. Proc. Natl. Acad. Sci. U. S. A. doi: 10.1073/pnas.0804598105.

Hattori, T., Müller, C., Gebhard, S., Bauer, E., Pausch, F., Schlund, B., et al. (2010). SOX9 is a major negative regulator of cartilage vascularization, bone marrow formation and endochondral ossification. Development. doi: 10.1242/dev.045203.

Hines, E. A., Jones, M. K. N., Verheyden, J. M., Harvey, J. F., and Sun, X. (2013). Establishment of smooth muscle and cartilage juxtaposition in the developing mouse upper airways. Proc. Natl. Acad. Sci. U. S. A. doi: 10.1073/pnas.1313223110.

Hirashima, T., and Adachi, T. (2015). Procedures for the Quantification of Whole-Tissue Immunofluorescence Images Obtained at Single-Cell Resolution during Murine Tubular Organ Development. PLoS One 10, e0135343. doi: 10.1371/journal.pone.0135343.

Kishimoto, K., Furukawa, K., LuzMadrigal, A., Yamaoka, A., Matsuoka, C., Habu, M., et al. (2019). Induction of tracheal mesoderm and chondrocyte from pluripotent stem cells in mouse and human. bioRxiv.

Kishimoto, K., Tamura, M., Nishita, M., Minami, Y., Yamaoka, A., Abe, T., et al. (2018). Synchronized mesenchymal cell polarization and differentiation shape the formation of the murine trachea and esophagus. Nat. Commun. doi: 10.1038/s41467-018-05189-2.

Komatsu, N., Aoki, K., Yamada, M., Yukinaga, H., Fujita, Y., Kamioka, Y., et al. (2011). Development of an optimized backbone of FRET biosensors for kinases and GTPases. Mol. Biol. Cell 22, 4647–56. doi: 10.1091/mbc.E11-01-0072.

Komatsu, N., Terai, K., Imanishi, A., Kamioka, Y., Sumiyama, K., Jin, T., et al. (2018). A platform of BRET-FRET hybrid biosensors for optogenetics, chemical screening, and in vivo imaging. Sci. Rep. doi: 10.1038/s41598-018-27174-x.

Lefebvre, V., Angelozzi, M., and Haseeb, A. (2019). SOX9 in cartilage development and disease. Curr. Opin. Cell Biol. doi: 10.1016/j.ceb.2019.07.008.

Mammoto, T., Mammoto, A., Torisawa, Y. suke, Tat, T., Gibbs, A., Derda, R., et al. (2011). Mechanochemical Control of Mesenchymal Condensation and Embryonic Tooth Organ Formation. Dev. Cell. doi: 10.1016/j.devcel.2011.07.006.

Mangan, S., and Alon, U. (2003). Structure and function of the feed-forward loop network motif. Proc. Natl. Acad. Sci. U. S. A. doi: 10.1073/pnas.2133841100.

Mason, J. M., Morrison, D. J., Basson, M. A., and Licht, J. D. (2006). Sprouty proteins: Multifaceted negative-feedback regulators of receptor tyrosine kinase signaling. Trends Cell Biol. doi: 10.1016/j.tcb.2005.11.004.

Murakami, S., Kan, M., McKeehan, W. L., and De Crombrugghe, B. (2000). Up-regulation of the chondrogenic Sox9 gene by fibroblast growth factors is mediated by the mitogen-activated protein kinase pathway. Proc. Natl. Acad. Sci. U. S. A. doi: 10.1073/pnas.97.3.1113.

Nakamura, Y., Yamamoto, K., He, X., Otsuki, B., Kim, Y., Murao, H., et al. (2011). Wwp2 is essential for palatogenesis mediated by the interaction between Sox9 and mediator subunit 25. Nat. Commun. doi: 10.1038/ncomms1242.

Nel-Themaat, L., Vadakkan, T. J., Wang, Y., Dickinson, M. E., Akiyama, H., and Behringer, R. R. (2009). Morphometric analysis of testis cord formation in sox9-egfp Mice. Dev. Dyn. doi: 10.1002/dvdy.21954.

Oh, C.-D., Chang, S.-H., Yoon, Y.-M., Lee, S.-J., Lee, Y.-S., Kang, S.-S., et al. (2000). Opposing Role of Mitogen-activated Protein Kinase Subtypes, Erk-1/2 and p38, in the Regulation of Chondrogenesis of Mesenchymes. J. Biol. Chem. 275, 5613–5619. doi: 10.1074/jbc.275.8.5613.

Park, J., Zhang, J. J. R., Moro, A., Kushida, M., Wegner, M., and Kim, P. C. W. (2010). Regulation of Sox9 by Sonic Hedgehog (Shh) is essential for patterning and formation of tracheal cartilage. Dev. Dyn. doi: 10.1002/dvdy.22192.

Sala, F. G., Del Moral, P. M., Tiozzo, C., Al Alam, D., Warburton, D., Grikscheit, T., et al. (2011). FGF10 controls the patterning of the tracheal cartilage rings via Shh. Development. doi: 10.1242/dev.051680.

Shen-Orr, S. S., Milo, R., Mangan, S., and Alon, U. (2002). Network motifs in the transcriptional regulation network of Escherichia coli. Nat. Genet. doi: 10.1038/ng881.

Shyer, A. E., Rodrigues, A. R., Schroeder, G. G., Kassianidou, E., Kumar, S., and Harland, R. M. (2017). Emergent cellular self-organization and mechanosensation initiate follicle pattern in the avian skin. Science (80-.). doi: 10.1126/science.aai7868.

Snowball, J., Ambalavanan, M., Whitsett, J., and Sinner, D. (2015). “Endodermal Wnt signaling is required for tracheal cartilage formation.” Dev. Biol. doi: 10.1016/j.ydbio.2015.06.009.

Tiozzo, C., Langhe, S. De, Carraro, G., Alam, D. Al, Nagy, A., Wigfall, C., et al. (2009). Fibroblast growth factor 10 plays a causative role in the tracheal cartilage defects in a mouse model of apert syndrome. Pediatr. Res. doi: 10.1203/PDR.0b013e3181b45580.

Turcatel, G., Rubin, N., Menke, D. B., Martin, G., Shi, W., and Warburton, D. (2013). Lung mesenchymal expression of Sox9 plays a critical role in tracheal development. BMC Biol. doi: 10.1186/1741-7007-11-117.

Yin, W., Kim, H. T., Wang, S., Gunawan, F., Wang, L., Kishimoto, K., et al. (2018). The potassium channel KCNJ13 is essential for smooth muscle cytoskeletal organization during mouse tracheal tubulogenesis. Nat. Commun. doi: 10.1038/s41467-018-05043-5.

